# L-citrulline ameliorates pathophysiology in a rat model of superimposed preeclampsia

**DOI:** 10.1101/2021.08.24.457514

**Authors:** Andy W. C. Man, Yawen Zhou, Uyen D. P. Lam, Gisela Reifenberg, Anke Werner, Alice Habermeier, Ellen I. Closs, Andreas Daiber, Thomas Münzel, Ning Xia, Huige Li

## Abstract

Preeclampsia, characterized by hypertension, proteinuria, and fetal growth restriction, is one of the leading causes of maternal and perinatal mortality. By far, there is no effective pharmacological therapy for preeclampsia. The present study was conducted to investigate the effects of L-citrulline supplementation in Dahl salt-sensitive rat, a model of superimposed preeclampsia. Parental DSSR were treated with L-citrulline (2.5 g/L in drinking water) from the day of mating to the end of lactation period. Blood pressure of the rats was monitored throughout pregnancy and markers of preeclampsia were assessed. Endothelial function of the pregnant DSSR was assessed by wire myograph. L-citrulline supplementation significantly reduced gestational hypertension, proteinuria, and levels of circulating soluble fms-like tyrosine kinase 1 in DSSR. L-citrulline improved maternal endothelial function by augmenting the production of nitric oxide in the aorta and improving endothelium-derived hyperpolarizing factor-mediated vasorelaxation in resistance arteries. L-citrulline supplementation improved placental insufficiency and fetal growth, which were associated with an enhancement of angiogenesis and reduction of fibrosis and senescence in the placentas. In addition, L-citrulline downregulated genes involved in the toll-like receptor 4 and nuclear factor-κB signaling pathway. In conclusion, this study shows that L-citrulline supplementation reduces gestational hypertension, improves placentation and fetal growth in a rat model of superimposed preeclampsia. L-citrulline supplementation may represent an effective and safe therapeutic strategy for preeclampsia that benefit both the mother and the fetus.

## Introduction

Preeclampsia is a gestation complication affecting 2%to 8%of all pregnancies worldwide and is a leading cause of maternal and perinatal mortality ^1^. Preeclampsia commonly occurs during the second half of pregnancy, and is characterized by hypertension, proteinuria, maternal organ damages and fetal growth restriction ^2^. Preeclampsia can leave long-term metabolic and cardiovascular risk to both mother and children. Women with a history of preeclampsia have approximately 2-fold increased risk of developing cardiovascular diseases and around 10-fold increased risk of chronic kidney diseases ^3^. Fetal growth restriction limits the growth potential of the offspring and increases the risk of diseases later at adult age ^4^.

The origin of preeclampsia is unclear. Accumulating studies have evidenced the involvement of multifactorial mechanisms including the imbalance in angiogenic factors, aberrant inflammatory response, increased placental oxidative stress and placental aging ^5, 6^. The production of anti-angiogenic factors including soluble fms-like tyrosine kinase 1 (sFIt-1) from the ischemic placentas can cause endothelial dysfunction, intravascular inflammation and activation of the hemostatic system ^7^. Due to the complex etiology of preeclampsia and safety concerns on drug usage during pregnancy, there is still no effective pharmacological treatments available for preeclampsia ^1^. An ideal therapeutic should have protective effects in lowering blood pressure, ameliorate maternal phenotypes and improving fetal growth and survival.

Preeclampsia can be considered as a vascular disorder. A functional and adequately vascularized placenta is crucial for a healthy pregnancy and birth outcome ^8^. Placental dysfunction results in a decrease in the angiogenic vascular endothelial growth factor (VEGF) and placental growth factor (PlGF) and the release of deleterious placental factors including sFIt1 into the maternal circulation causing generalized endothelial dysfunction ^9, 10^. Nitric oxide (NO) donors have potent vasodilator effect and have been shown to improve blood flow in the fetoplacental circulation of pregnancies affected by mild preeclampsia ^11^.

Several animal studies and clinical trials have tested the effect of L-arginine, a semi-essential amino acid and the substrate for vascular NO formation, in treating preeclampsia ^12–14^. Nevertheless, the bioavailability of arginine is relatively low, as 60%of oral arginine evades intestinal metabolism and another 9%is metabolized by liver ^15^. Citrulline is the endogenous precursor to arginine. Compared to arginine, citrulline is more effective to augment NO as it can bypass hepatic metabolism, and is not metabolized by arginase ^15^. Some studies suggested that the protective effect of citrulline treatment may not analogous with L-arginine treatment ^16^, and the two amino acids differentially regulate gene expression ^17^. Moreover, citrulline has been shown to possess protein anabolic effect, which increases nitrogen balance in rats ^18^ and enhances muscle protein synthesis in human ^19^ more efficiently than arginine. In addition, usage of citrulline has an even greater safety profile than that of arginine, as none of the trials reported any adverse effects ^20^. Citrulline may therefore be more effective in the treatment of pre-eclampsia. Maternal citrulline supplementation in a rat model of low protein diet-induced intrauterine growth restriction (IUGR) has been shown to enhance fetal growth and protein metabolism ^21^. Nevertheless, the effects of maternal citrulline supplementation in targeting preeclampsia remain unclear.

Dahl sat sensitive rats (DSSR) is a genetic model of salt-induced hypertension, renal injury and insulin resistance ^22^. Recent studies have reported the preeclamptic phenotype and fetal growth restriction in DSSR, which reminisce many characteristics of preeclampsia in human ^23–27^. During pregnancy, DSSR spontaneously displays increased blood pressure, proteinuria, placental hypoxia and reduced uteroplacental blood flow. These phenotypes are associated with increased placental production of tumour necrosis factor-α (TNF-α), hypoxia-inducible factor-1α (HIF-1α), and sFlt-1 ^23^. In the present study, we evaluated whether citrulline could alleviate the pathophysiology of preeclampsia. We monitored blood pressure, evaluated the level of preeclampsia markers, placenta phenotypes, vascular function, and pregnancy outcomes in pregnant DSSR with or without citrulline supplementation.

## Method

### Animals

DSSR were from Charles River Laboratories (Sulzfeld, Germany). F0 parental DSSR were fed with standard chow diet *ad libitum*. Mating was set at the age of 12 weeks old. Pregnancy was confirmed by checking the plug. Only the females giving the first birth were used in the study. Female rats from the same litter were randomized to receive either normal water or L-citrulline (2.5 g/L in drinking water) from the day of mating to the end of lactation period. Citrulline were replaced every 2 days during the treatment period. Female rats from both groups were studied either during pregnancy (GD 12 = 9 rats per group; GD 21 = 10 rats per group) or postpartum (control = 4 cohort, citrulline = 5 cohort). At the terminal day of experiments, isoflurane and intraperitoneal injection of pentobarbital were used for euthanasia. For GD12 experiments, pregnant rats were sacrificed at GD12 to measure the litter size and embryo weight, to collect aorta and mesenteric arteries for *ex vivo* vascular function experiments. For GD21 experiments, blood pressure of the pregnant rats was measured during each week of pregnancy. Rats were sacrificed at GD21 to measure the litter size and embryo weight, to collect urine and serum samples, and to collect umbilical vein for *ex vivo* vascular function experiments and placenta samples for Western blotting, qPCR and staining. For postpartum experiments, the body weights of the pup were followed till day 8. The animal experiment was approved by the responsible regulatory authority (Landesuntersuchungsamt Rheinland-Pfalz; 23 177-07/G 16-1-038) and was conducted in accordance with the German animal protection law and the National Institutes of Health (NIH) Guide for the Care and Use of Laboratory Animals.

### Blood pressure measurement

Systolic, diastolic, and mean blood pressure were measured noninvasively in conscious rats (control = 6, citrulline = 7) by a computerized system (CODA Monitor, Kent Scientific) as described in our previous studies ^28–30^. Rats were restrained in individual holders. The occlusion cuff and the volume-pressure recording cuff were placed close to the base of the tail. After an adaptation period of 30 minutes on a 37°C warm pad, 10 preliminary measurements were performed before actual measurement. Rats were acclimated for three consecutive days prior to the actual measurement. Results are presented as the mean of at least 15 recordings on each occasion taken. The measurements were performed at the same time of a day from 2 pm to 4 pm by the same investigator.

### Isometric tension studies

The vascular function of the second order mesenteric arteries and umbilical veins was accessed by wire myograph as described in our previous studies ^29^. Vessels were dissected free of adherent connective tissues and placed in cold modified Krebs-Ringer bicarbonate buffer under continuous aeration with 95%O2/5%CO2. Vessel rings with 2–3 mm long were suspended in the chambers of a Mulvany–Halpern wire myograph system (620M, Danish Myo Technology A/S, Aarhus, Denmark). Isometric force was recorded by a PowerLab/8SP system (AD Instruments Inc., Colorado Springs, CO, USA). The preparations were equilibrated for 30 mins at the optimal resting tension. The viability of the endothelium was tested by the relaxation response to a single dose of acetylcholine (10^−4^ M) after obtaining a reference contraction to 60 mM potassium chloride (KCl) twice prior to the actual experiment. For actual experiment, preparations were incubated for 30 mins with or without different inhibitors (10^−4^ M *N*^G^-nitro-L-arginine methyl ester, L-NAME and 10^−5^ M indomethacin, indo). Preparations were then pre-contracted by exposing to increasing concentrations of phenylephrine (PE, 10^−9^ to 10^−5^ M). Endothelium-dependent relaxation was examined by exposure to increasing concentrations of acetylcholine (10^−9^ to 10^−4^ M). Change in tension is expressed as percentage of the PE contraction (~80%of the reference KCl contraction). For umbilical veins, endothelium-dependent contraction was examined by exposure to increasing concentrations of acetylcholine (10^−9^ to 10^−4^ M). Area under the contraction curve (AUCC) was measured in different dose-dependent curve of the preparation with or with L-NAME. The difference between AUCC (ΔAUCC) was calculated to examine the L-NAME-induced contraction.

### Cell culture

Human umbilical vein endothelial cells (HUVEC)-derived EA.hy 926 cells were cultured in Dulbecco’s modified Eagle’s medium (DMEM; Sigma) with 10%fetal calf serum (FCS; PAA Laboratories, Germany), 2 mM L-glutamine, 2 mM sodium pyruvate, 1 %penicillin/streptomycin, and 1%HAT (hypoxanthine, aminopterin, and thymidine) (Life Technologies/Thermo Fisher Scientific). Cells were incubated in moisturized chamber with 10%CO2. Before experiments, confluent EA.hy 926 cells were cultured for 24 h in DMEM with 10%FCS, 2 mM L-glutamine, 2 mM sodium pyruvate, 1 %penicillin/streptomycin, and 1%HT (hypoxanthine and thymidine). Cells were incubated with or without 1 mM *N*^G^-nitro-L-arginine methyl ester (L-NAME) for 1 hour before treated with 2.5 mM L-citrulline for 24 hours. The cells were then collected for RNA extraction.

### Senescence assay

EA.hy 926 cells were cultured in DMEM with 10%FCS, 2 mM L-glutamine, 2 mM sodium pyruvate, 1 %penicillin/streptomycin, 1%HT and 5%serum from either control or citrulline-treated rats. After cultured for 3 days, cells were fixed and stained with senescence detection kit (ab65351, Abcam). Beta-galactosidase (SA-beta-Gal) activity is a known characteristic of senescent cells. The SA-beta-Gal is present only in senescent cells. In brief, the cells were fixed in fixative solution for 15 minutes and stained with SA-beta-gal solution at 37°C overnight. The development of the bluish-green color was observed using a light microscope. Experiment was repeated triplicate independently.

### Gene expression studies by quantitative PCR

Total RNA from randomly selected placentas from different rats and cell culture was isolated using peqGOLD TriFast™ (PEQLAB) and cDNA was reverse transcribed using High Capacity cDNA Reverse Transcription Kit (Applied Biosystems) according to our previous publication ^31^. QPCR was performed using SYBR Green JumpStart™ *Taq* Ready-Mix™ (Sigma-Aldrich) on an iCycler Real-Time PCR Detection System (Bio-Rad). Quantification was achieved by the difference in quantification cycles (ΔΔCt) values that were normalized to reference genes (GAPDH for rat samples or β-actin for HUVEC samples). Relative gene expression of target gene in each sample was expressed as the percentage of control. Specificity of the qPCR primers were checked by melting curve analysis or gel electrophoresis of the qPCR product. The sequence of the primers used are listed in Supplementary Table 1.

### Protein expression by Western blotting

Subset of randomly selected aorta and placenta samples were homogenized in RIPA buffer with 1% (v/v) proteinase inhibitor cocktail (#78442, Thermo Fisher Scientific, USA). Same amount of lysates protein was loaded and separated in SDS-PAGE. The resolved proteins were transferred onto nitrocellulose membranes and probed with specific primary antibody at 4°C overnight with agitation. GAPDH or β-actin was probed as a loading control. The protein bands were visualized using enhanced chemiluminescence (ECL) reagents (GE Healthcare, USA) and developed in Fujitsu Biomedical film (Fujitsu, Japan). Quantification protein expression was based on the ratio of target protein to GAPDH. The following primary antibodies were used: anti-CD31 (#PA5-16301, Invitrogen, 1:1000), anti-PlGF (#PA5-79814, Invitrogen, 1:1000), anti-VEGF (#MA1-16629, Invitrogen, 1:1000), anti-p-eNOS (#9571S, Cell Signaling, 1:1000), anti-eNOS (#610297, BD, 1:1000), anti-p-VASP (#3114S, New England Biolabs, 1:1000), anti-VASP (#3132S, New England Biolabs, 1:1000), anti-β-tubulin I (T7816, Sigma-Aldrich, 1:30000), and anti-GAPDH (2251-1, Epitomics, 1:30000).

### Masson’s Trichome Staining

Placentas were fixed in 4%paraformaldehyde and embedded in paraffin. Microtome sectioning was performed to obtain slides with thickness of 5 μm. After deparaffination and rehydration, Trichrome Stain Kit (Abcam, UK) was used to analyze collagen fibers according to manufacturer’s instructions. In brief, the slides were incubated in the pre-heated Bouin’s Solution at 60°C for 1 hour. The yellow color on the slides was removed by rinsing in running tap water followed by staining in Working Weigert’s Iron Hematoxylin Solution. The slides were then rinsed in deionized water and stained in Working Phosphomolybdic/Phosphotungstic Acid Solution followed by Aniline Blue Solution and 1%acetic acid. The slides were then rinsed, dehydrated, and mounted. Collagen staining per images was quantified by NIH ImageJ software. Staining was performed in 6 different placenta samples from each group.

### Immunohistochemistry (IHC) staining

Placentas were fixed in 4%paraformaldehyde and embedded in paraffin. Microtome sectioning was performed to obtain slides with thickness of 5 μm. After deparaffination and rehydration, slides were immersed in warm EnVision FLEX target Retrieval Solution (#K8004, Agilent, USA) for antigen retrieval. The slides were then immersed in peroxidase blocking solution to inhibit the activity of peroxidase. The sections were incubated with primary antibody of CD31 (#PA5-16301, Invitrogen, 1:50) in 4°C overnight. After washing twice in phosphate buffered saline with 0.1%Tween 20 (PBST), the sections were incubated with HRP-linked anti-rabbit antibody (#K4002, Agilent, USA) in room temperature for 1 hour. After washing twice in PBST, the sections were incubated with DAB (3,3’-Diaminobenzidine) staining kit (#K3468, Agilent, US) for 5 minutes at room temperature. The slides were rinsed in distilled water, followed by dehydration and mounting.

### ELISA assay

Serum level of rat soluble fms-like tyrosine kinase-1 (sFlt-1, #MBS2602003) and placenta growth factor (PIGF, #MBS026910) were examined using enzyme-linked immunosorbent assay (ELISA) according to the manufacturer’s instruction (Mybiosource, USA). In brief, 100 μl serum were added as samples. Samples and standard were then incubated at room temperature for 90 minutes with gentle shaking. After washing twice, biotinylated antibodies were added and incubated at room temperature for another 60 minutes. After washing for three times, HRP-avidin was added and incubated for 30 minutes at room temperature with gentle shaking. After washing for five times color reagent was added in dark with gentle shaking. After 30 minutes of incubation, stop solution was added and the absorption was read at 450 nm immediately with Sunrise™ microplate reader (Tecan Group, Switzerland) and analyzed by Magellan™ software (Tecan Group, Switzerland).

### Urine Chemistry Detection

Urine samples were collected directly from the bladder after euthanasia. Urine creatinine was measured by a blood chemical analyzer (Reflotron, Roche Co., Germany) using the specified analysis kits supplied from the manufacturer. Urine protein was measured by bicinchoninic acid protein assay.

### *In vitro* NO production assays

Electron paramagnetic resonance (EPR)-based NO-trapping technique with iron-diethyldithiocarbamate (Fe(DETC)2) colloid was used to assess the NO production in the aortas *in vitro* as previously described ^32–34^. Rat aorta was cut into rings of 3 mm in length. The samples were incubated with 10 μM calcium ionophore (A23187) and 200 μM Fe(DETC)2 colloid solution at 37 °C for 60 min in Krebs-HEPES buffer. EPR conditions: B0= 3276 G, sweep=115 G, sweep time=60 s, modulation=7000 mG, MW power=10 mW, gain=9×10^2^ using a Miniscope MS400 from Magnettech (Germany).

### Statistics

Results were expressed as mean ± SEM (standard error of the mean). Student’s t test was used for comparison between two groups or one-way ANOVA was used for comparison between multiple groups. *P* values < 0.05 were considered significant. GraphPad Prism (GraphPad Software, La Jolla, CA, USA) was used to generate graphs and statistical analysis.

## Results

### Citrulline reduces preeclamptic phenotypes during pregnancy in DSSR

We investigated the effect of maternal L-citrulline supplementation in the pregnant DSSR. Blood pressure was monitored from the day of mating to the third week of pregnancy. Blood pressure was similar in the first week of pregnancy among control and citrulline-treated DSSR. Citrulline supplementation reduced the systolic blood pressure (SBP) starting from the first week of pregnancy **(Figure 1A)**. There was a significant reduction of the diastolic blood pressure (DBP) in the second and third week of pregnancy in citrulline-treated DSSR **(Figure 1B)**. Citrulline supplementation resulted in a significant decrease in mean arterial pressure (MAP) beginning from the second weeks and the effect sustained during the late pregnancy **(Figure 1C)**.

**Fig 1.**
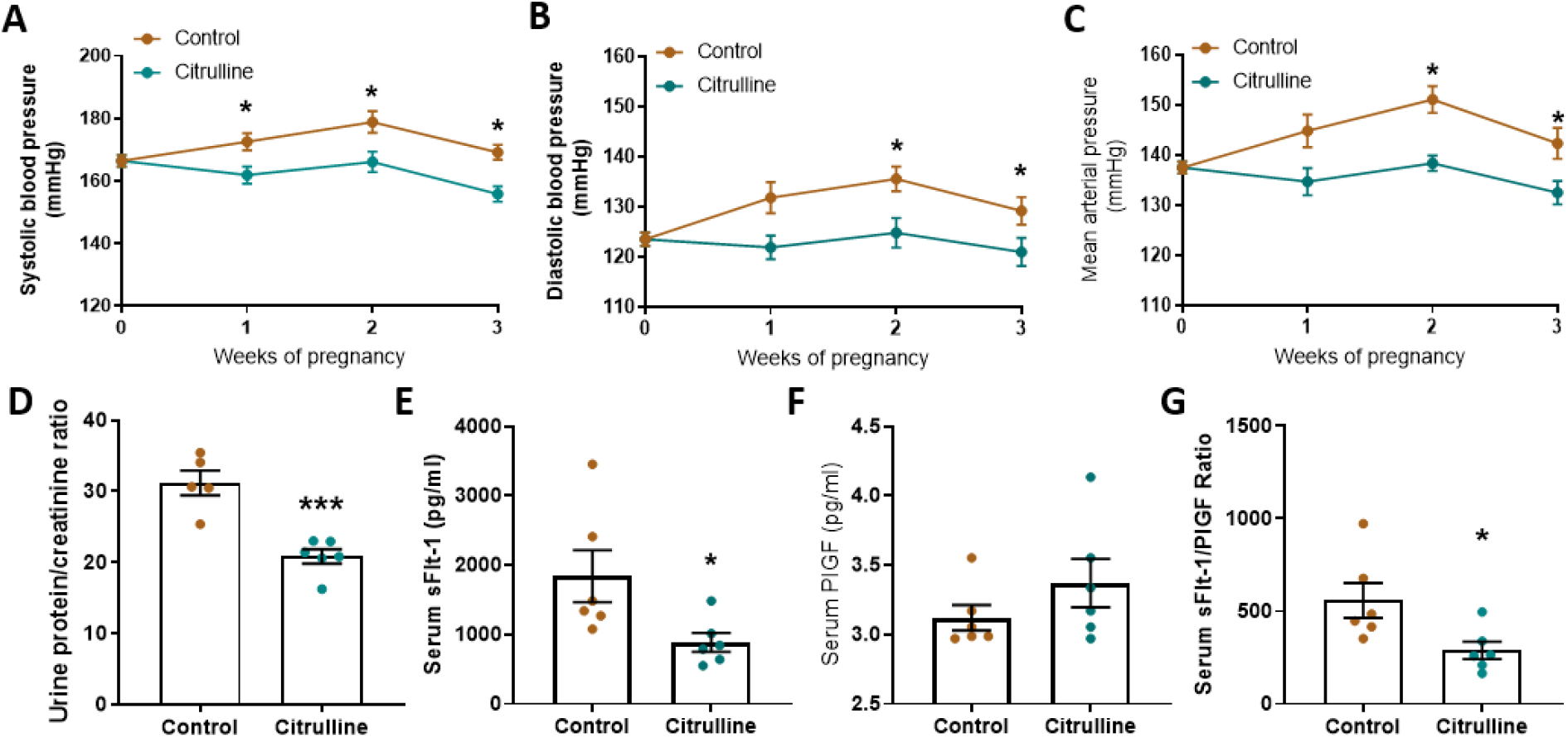
Citrulline reduces preeclamptic phenotypes in DSSR. L-citrulline (2.5g/L water) was administrated from the day of mating to the end of the experiments. Systolic blood pressure **(A)**, diastolic blood pressure **(B)**, and mean arterial pressure **(C)** were measured by tail-cuff method in pregnant DSSR. Control = 6, citrulline = 7. The experiment was terminated at gestational day (GD) 21. The urine protein content was measured by bicinchoninic acid assay and creatinine level was measured by a chemical analyser. The urine protein-to-creatinine ratio was calculated **(D)**. Levels of soluble fms-like tyrosine kinase-1 (sFlt-1) (**E)** and placental growth factor (PlGF) **(F)** were measured in serum collected at GD 21 with specific enzyme-linked immunosorbent assay (ELISA) kit. The serum sFlt-1-to-PlGF ratio was calculated as a preeclampsia marker **(G)**. *, p<0.05, ***, p<0.001. Data are presented as mean ± SEM.

It is known that DSSR exhibits an increase in proteinuria over the course of pregnancy ^23^. Urine protein-to-creatinine ratio was significantly reduced by citrulline supplementation **(Figure 1D)**. Circulating level of sFlt-1 was significantly reduced in citrulline-treated DSSR **(Figure 1E)**, whereas serum level of PlGF had no significant difference between control and citrulline-treated DSSR **(Figure 1F)**. Citrulline supplementation reduced the serum sFlt-1/PIGF ratio at late pregnancy of DSSR **(Figure 1G)**.

### Citrulline improves placental insufficiency during pregnancy and improves fetal growth in DSSR

Placental insufficiency causes intrauterine growth restriction during preeclampsia, which leads to the reduction of fetus size in both human patients and animal models ^35^. Citrulline supplementation significantly improved fetal growth in DSSR. At gestation day 12, pregnant DSSR were sacrificed to study the intrauterine growth. The number of embryos per litter was not significantly different in the control and citrulline-treated DSSR **(Figure 2A)**. Citrulline-treated DSSR had significantly larger embryos, as measured by increased weight **(Figure 2B)** and size **(Figure 2C)**. At gestation day 21, the number of fetuses per litter was not significantly different between control and citrulline-treated DSSR **(Figure 2D)**. Placental efficiency, as quantified by the ratio of pup-to-placenta weight, was improved by citrulline supplementation **(Figure 2E)**. A high placenta weight-to-pup weight ratio is usually associated with growth restriction and represents a reduced nutrient transport capacity of the placentas. The birth weight of the offspring was significantly increased in the citrulline-treat DSSR **(Figure 2F and G)**.

**Fig 2.**
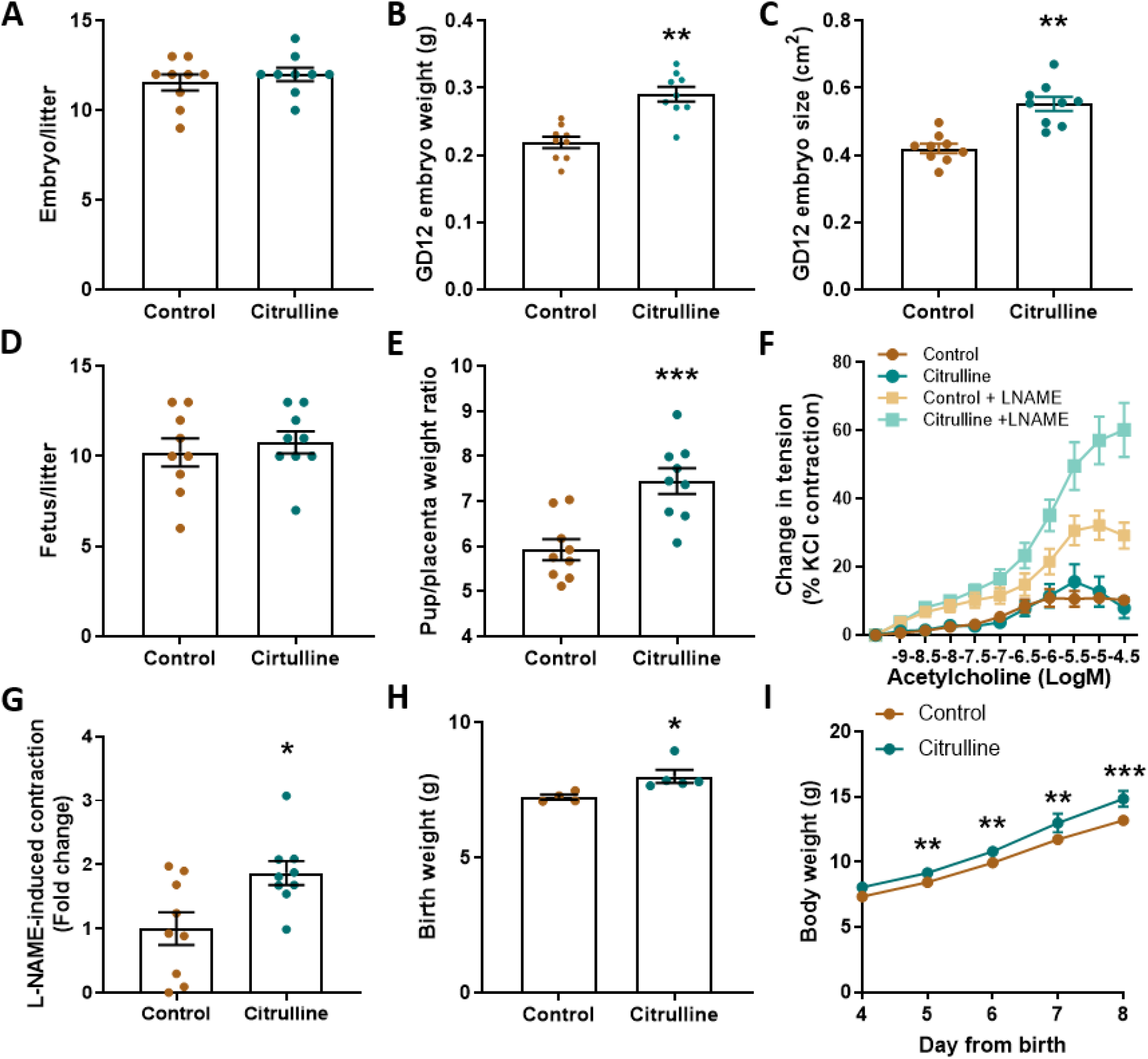
Citrulline improves placental insufficiency and improves fetal growth in DSSR. L-citrulline (2.5 g/L in drinking water) was administrated from the day of mating to the end of the experiments. At gestational day (GD) 12, the number of embryos per litter **(A)**, and the weight **(B)** and size **(C)** of the embryos were measured. At gestational day (GD) 21, the number of fetuses per litter **(D)** and the ratio of pup-to-placenta weight **(E)** were measured. The vascular responsiveness of the GD 21 umbilical vein was measured using wire myography. Preparations were incubated for 30 mins with or without NG-nitro-l-arginine (L-NAME, 10^−4^ M). Preparations were then exposed to increasing concentrations of acetylcholine to induce contraction **(F)**. L-NNA-induced contraction was calculated by the difference between area under the contraction curve (ΔAUCC) **(G)**. The birth weight of the offspring was measured (**H)** and the body weight of the rats was measured from the age of 4 to 8-day **(I)**. *, p<0.05, **, p<0.01, ***, p<0.001. Data are presented as mean ± SEM.

The umbilical vessels provide fetal blood supply, and preeclampsia may lead to reduced placental perfusion and impaired fetal development. Response to acetylcholine was studied in the umbilical vein using in a wire myograph system. Acetylcholine induced contraction of the umbilical vein (**Figure 2H**), which is consistent with previous reports ^36, 37^. In the presence of NOS inhibitor L-NAME, acetylcholine-induced contraction was much larger in the umbilical vein from citrulline-treated DSSR (**Figure 2I**), indicative of improved NO production by citrulline. The L-NAME-induced contraction was used as an indicator of endogenous NO production that counterbalances vasoconstriction ^38^.

### Citrulline improves endothelial function in pregnant DSSR

EPR spectra of NO-Fe(DETC)2 were obtained and used to assess the *in vitro* NO production in the aorta of DSSR **(Figure 3A)**. Quantification result suggested a significantly higher NO production in the aorta of citrulline-treated DSSR compared to control **(Figure 3B)**. Also, citrulline supplementation augmented endothelial NO signaling in DSSR. The total protein expression of both endothelial nitric oxide synthase (eNOS) and its key downstream target vasodilator-stimulated phosphoprotein (VASP) was significantly upregulated in the citrulline-treated DSSR **(Figure 3C)**. Relative phosphorylation of VASP was also significantly increased in the citrulline-treated DSSR.

**Fig 3.**
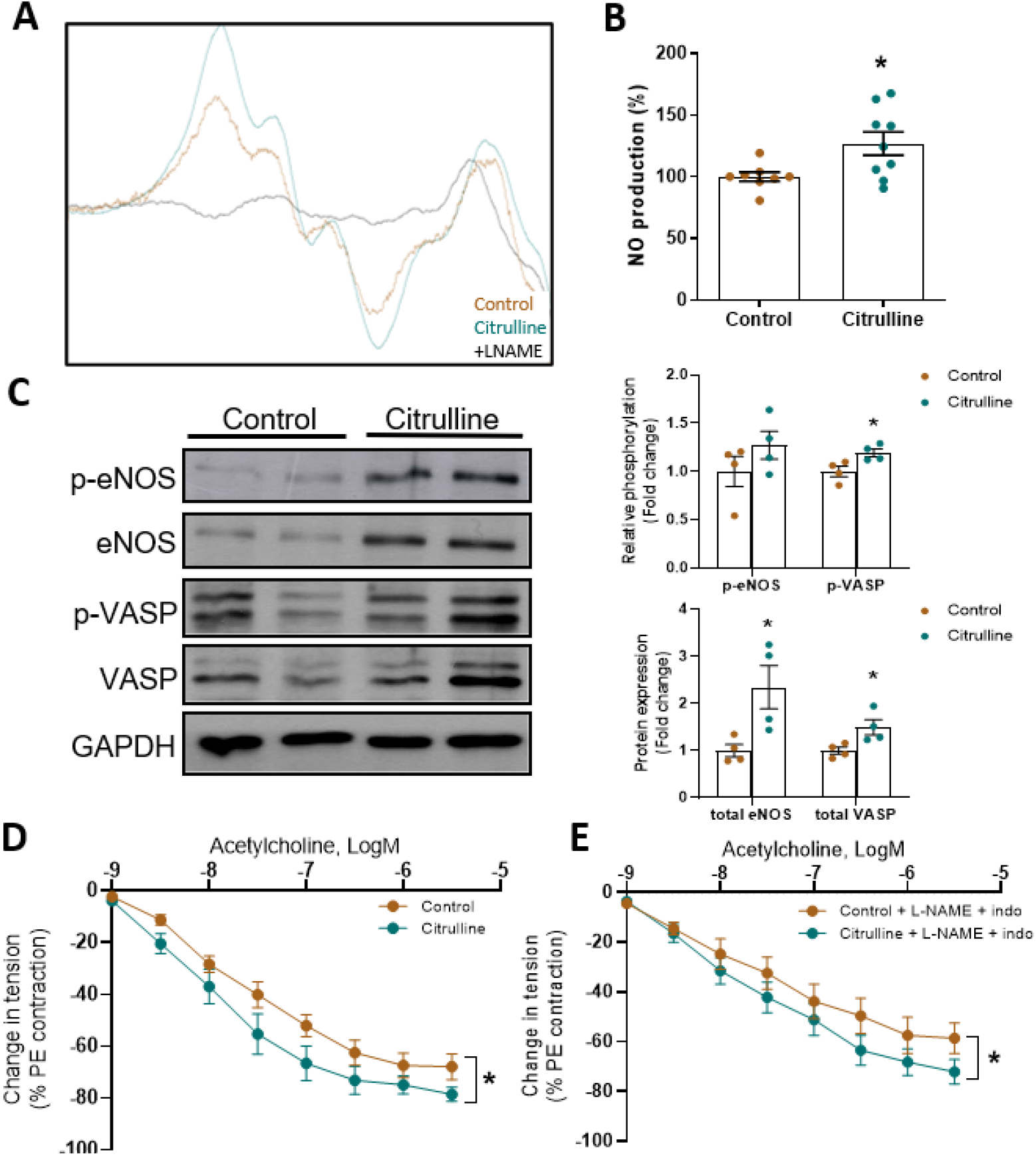
Citrulline improves endothelial function in pregnant DSSR. L-citrulline (2.5 g/L in drinking water) was administrated from the day of mating to the end of the experiments. At gestational day (GD) 12, nitric oxide (NO) production in the aorta of the pregnant rats was measured by electron paramagnetic resonance spectrometry (EPR) **(A)** and the relative percentage of the NO production was calculated **(B)**. Protein expression of phosphorylated and total eNOS, phosphorylated and total VASP in the aorta of the pregnant rats was analyzed by Western blotting. GAPDH was used as the internal control. Total protein level was normalized to internal control. Relative phosphorylation was normalized to total protein level **(C)**. The vascular responsiveness of the second order mesenteric artery of the pregnant rats was studied using wire myography. Preparations were pre-contracted with phenylephrine (PE). Basal endothelium-dependent relaxation was assessed by exposing the preparations to increasing concentrations of acetylcholine **(D)**. Endothelium-derived hyperpolarizing factor (EDHF) was assessed by exposing the preparations to acetylcholine in the presence of NG-nitro-l-arginine (L-NAME, 10^−4^ M) and indomethacin (indo, 10^−5^ M) **(E)**. Control =11 vessels, citrulline = 14 vessels; *, p<0.05, ***. Data are presented as mean ± SEM.

Citrulline supplementation significantly improved the vascular function of the second order mesenteric arteries in pregnant DSSR. Endothelium-dependent acetylcholine-mediated vasorelaxation was increased in citrulline-treated DSSR **(Figure 3D)**. Endothelium-derived hyperpolarizing factor (EDHF) was measured as the NO/prostaglandin-independent component of endothelium-dependent relaxation. In the presence of both L-NAME and indomethacin, citrulline significantly improved the EDHF-mediated relaxation of mesenteric arteries **(Figure 3E)**.

### Citrulline ameliorates fibrosis and promotes angiogenesis in the placenta

Histologic features of placentas in preeclamptic patients are characterized by chronic inflammation and fibrosis, while placental fibrosis could be triggered by transformation growth factor-β1 (TGF-β1) signaling ^39^. Therefore, we examined the histologic features of placentas in DSSR. Citrulline ameliorated placental fibrosis in DSSR. Collagen deposition in the placenta, as indicated by the blue color stained by Masson’s’ Trichome, was significantly reduced by citrulline **(Figure 4A and B)**. No significant changes of TGF-β1 expression was found. But the expression fibrotic markers, metalloproteinase 9 (MMP9) was downregulated in the placentas of citrulline-treated DSSR **(Figure 4C)**.

**Fig 4.**
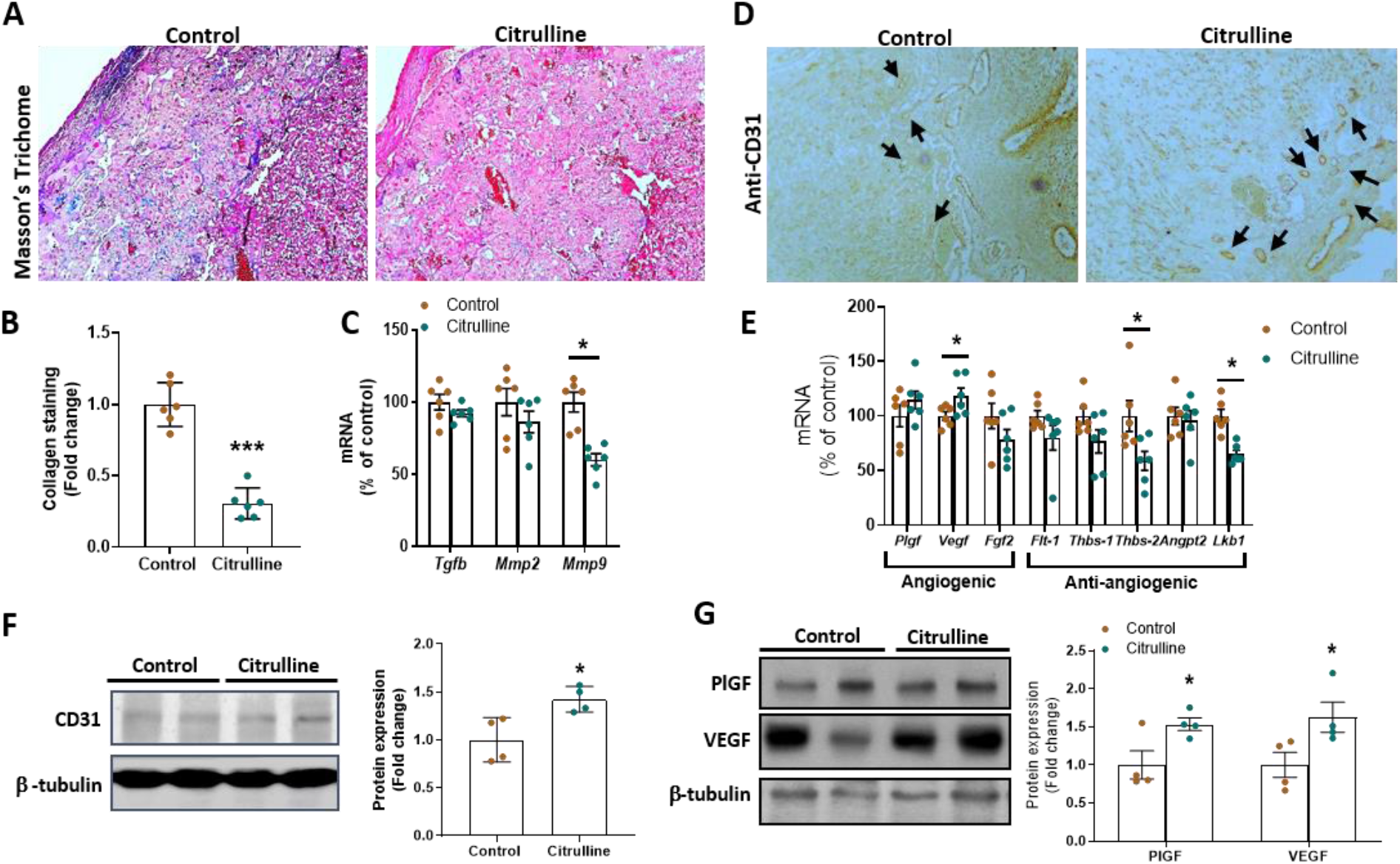
Citrulline ameliorates fibrosis and promotes angiogenesis in the placenta. L-citrulline (2.5 g/L in drinking water) was administrated from the day of mating to the end of the experiments. At gestation day (GD) 21, the placentas were fixed in formalin and embedded in paraffin. Placenta sections were staining by Masson’s Trichome staining kit and the blue colour indicates positive stain for collagen **(A)**. Collagen staining was quantified using ImageJ software **(B)**. n = 6 animals per group. The expression of remodelling genes including transforming growth factor beta (*Tgfb*) and matrix metalloproteinase 2 and 9 (*Mmp2* and *Mmp9*) in the placentas was analyzed by quantitative PCR **(C)**. The expression of endothelial cell marker CD31 was examined by immunohistochemistry (IHC) staining using anti-CD31 antibody **(D)**. Black arrows indicate small vessels with positive stain. The expression of angiogenic and anti-angiogenic genes in the placentas was measured by quantitative PCR **(E)**. The protein expression of CD31 **(F)**, placental growth factor (PlGF) and vascular endothelial growth factor (VEGF) **(G)** in the placentas was measured by Western blotting. β-tubulin was used as internal control. *, p<0.05. Data are presented as mean ± SEM.

Immunostaining of CD31, a marker of endothelial cells, was used to examine the placental vascularization. The results showed that the microvessels count was increased in the citrulline-treated DSSR placenta compared to control **(Figure 4D)**. Protein expression of CD31 was also increased in the citrulline-treated DSSR compared to control **(Figure 4F)**. Gene expression of anti-angiogenic factors was reduced, while the gene expression of angiogenic factors was increased in the placenta of citrulline-treated DSSR compared to control **(Figure 4E)**. In addition, the protein expression of PlGF and VEGF was upregulated in the citrulline-treated DSSR **(Figure 4G)**. These results suggested citrulline could improve angiogenesis in the placentas of DSSR.

### Citrulline ameliorates placental senescence during pregnancy in DSSR

Placental senescence has been associated with preeclampsia ^40^. Liver kinase B1 (LKB1) is a serine/threonine kinase that is highly expressed in senescent cells. LKB1 induces vascular senescence ^41^ and inhibits VEGF-induced angiogenesis ^42^. In the placentas of citrulline-treated DSSR, LKB1 was downregulated **(Figure 4E)**. Therefore, we further investigated the placental senescence in DSSR.

Gene expression of senescence markers (p16, p21, and p53) was downregulated in the placentas of citrulline-treated DSSR **(Figure 5A)**. Next, we examined whether the placental senescence was facilitated by circulating factors in the serum. Endothelial cells were incubated with the serum of pregnant DSSR. The results suggested that endothelial cells incubated with control serum had more positively β-gal-stained cells than those incubated with serum from citrulline-treated DSSR. **(Figure 5B)**. Gene expression of p53 in endothelial cells was significantly upregulated by control DSSR serum, but not by serum from citrulline-treated DSSR **(Figure 5C)**. These suggested citrulline supplementation may ameliorate placental senescence in DSSR.

**Fig 5.**
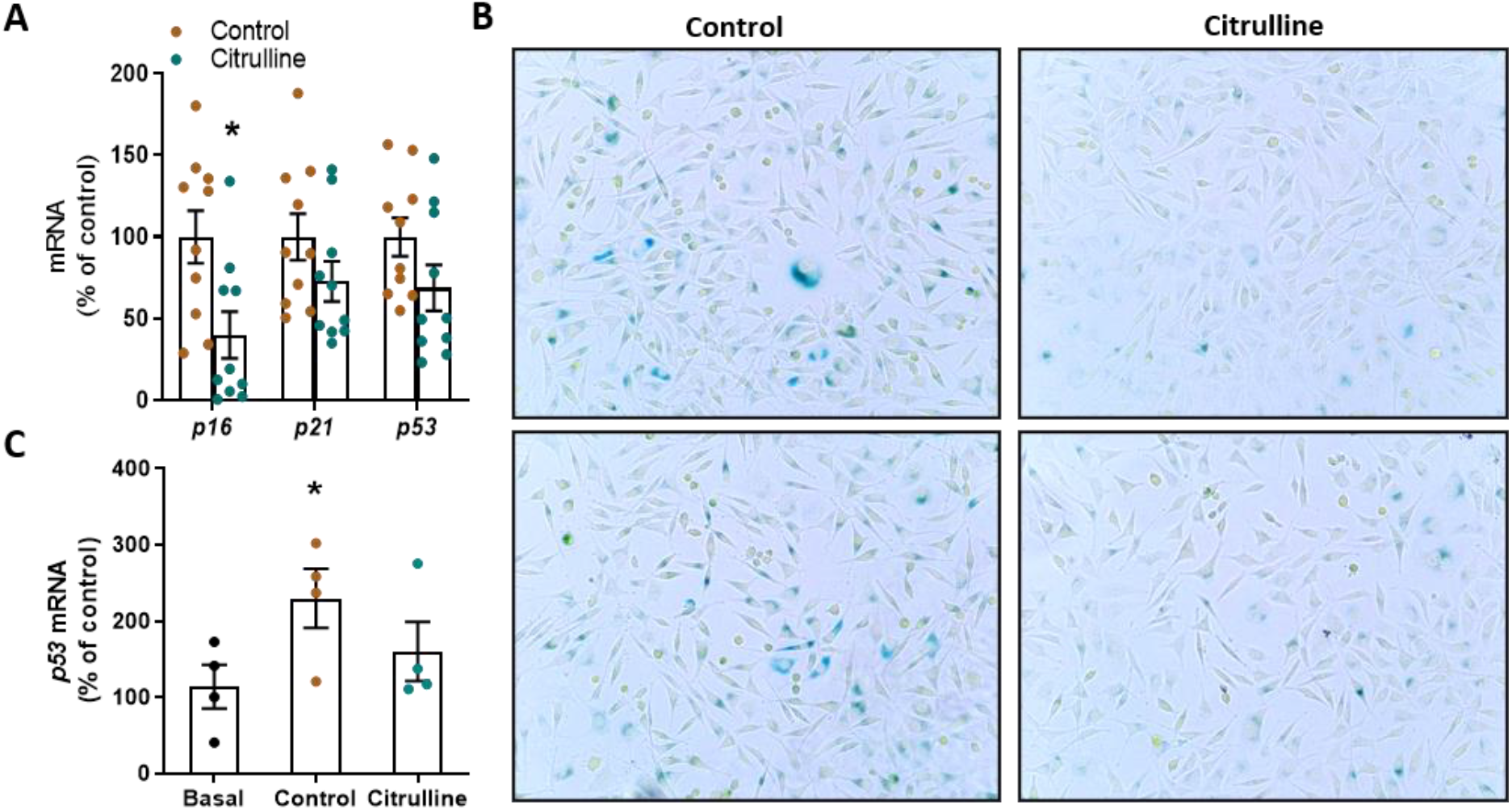
Citrulline ameliorates placenta senescence in DSSR. L-citrulline (2.5 g/L in drinking water) was administrated from the day of mating to the end of the experiments. At gestational day (GD) 21, the gene expression of senescence markers p16, p21 and p53 in the rat placentas was measured by qPCR **(A)**. Human umbilical vein endothelial cell line EA.hy926 were incubated either with the normal growth medium (basal), or additionally with serum (4 %) of control DSSR or citrulline-treated DSSR for 72 hours. Endothelial senescence was examined by β-galactosidase senescence detection kit (Abcam). The development of the bluish-green color indicates senescent cells **(B)**. Gene expression of p53 in the EA.hy926 incubated with the serum of either control or citrulline-treated rats was examined by qPCR **(C)**. *, p<0.05. Data are presented as mean ± SEM.

### Citrulline regulates differential gene expressions in DSSR placentas

High serum levels of toll-like receptor 4 (TLR4) and nuclear factor-κB (NF-κB) have been found in patients with preeclampsia, and are suggested to be used as biomarkers for predicting preeclampsia ^43^. Therefore, we studied the expression of genes involved in TLR4/NF-κB signaling in the placenta of DSSR. Gene expression levels of TLR4 and its downstream molecule myeloid differentiation factor 88 (MYD88), as well as the subunits of NF-kB (p65 and p50) in the DSSR placentas were significantly reduced by citrulline supplementation **(Figure 6A)**. Gene expression of downstream inflammatory markers of NF-κB, including inducible nitric oxide synthase (iNOS), vascular cell adhesion protein 1 (vcam-1) and intercellular adhesion molecule 1 (icam-1), was significantly downregulated in the placentas of citrulline-treated DSSR **(Figure 6B)**.

**Fig 6.**
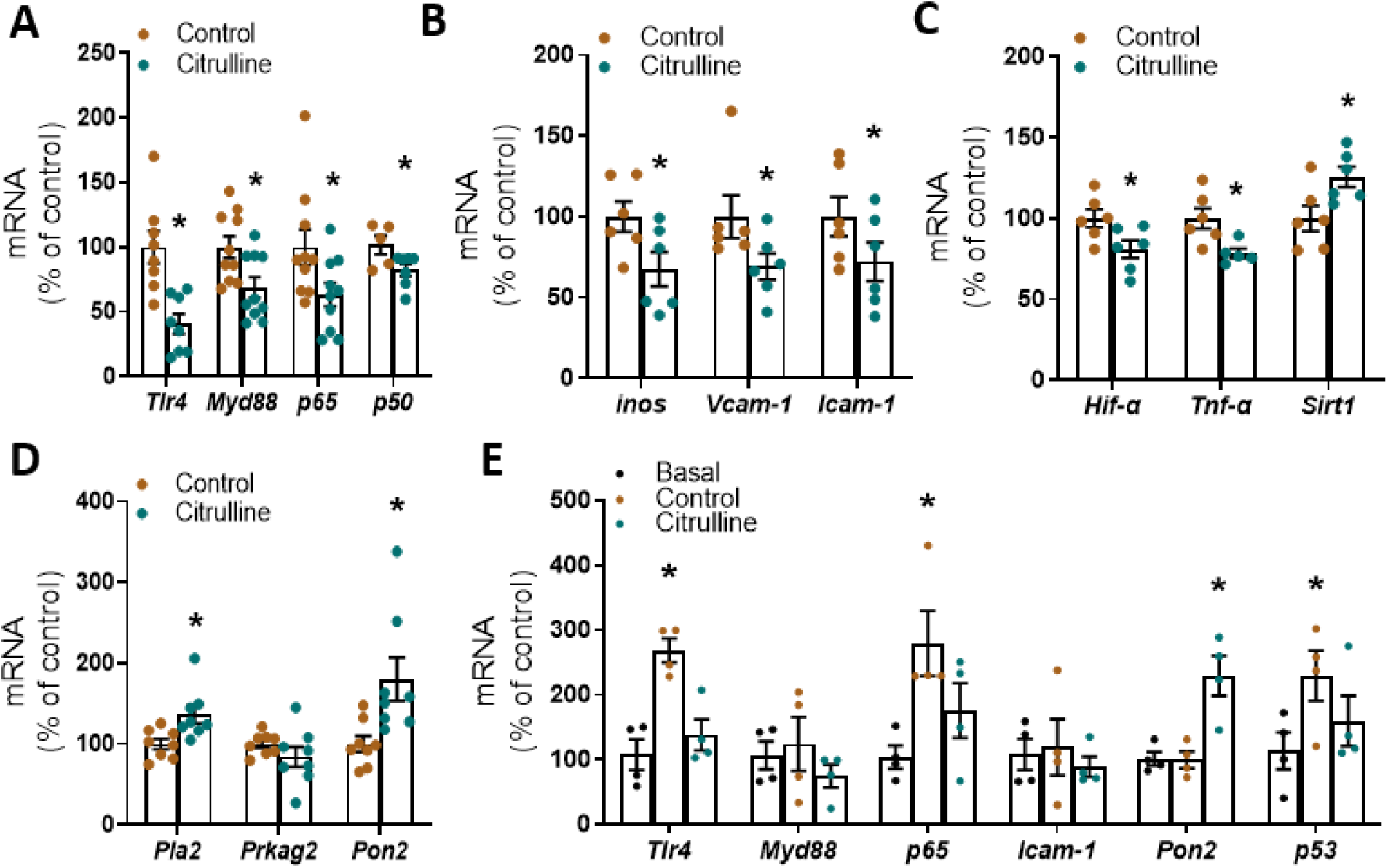
Citrulline regulates expression of different genes in DSSR placentas via NO-dependent or NO-independent mechanisms. L-citrulline (2.5 g/L in drinking water) was administrated from the day of mating to the end of the experiments. Placentas from gestational day 21 were collected for experiments. Expression of genes in the toll-like receptor 4 (TLR4) pathway including *Tlr4* and its downstream myeloid differentiation factor 88 (*Myd88*) and subunits of nuclear factor-kappa B (NF-κB) (*p65* and *p50*) in the DSSR placentas was measured by quantitative PCR **(A)**. Gene expression of downstream inflammation markers of NF-κB including inducible nitric oxide synthase (*inos*), vascular cell adhesion protein 1 (*Vcam-1*) and intercellular adhesion molecule 1 (*Icam-1*) in DSSR placentas was measured by quantitative PCR **(B)**. Gene expression of hypoxia inducible factor alpha (*Hif-α*), tumour necrosis factor alpha (*Tnf-α*), sirtuin 1 (*Sirt1*) in DSSR placentas was measured by quantitative PCR **(C)**. Expression of genes related to lipid metabolism, including protein kinase AMP-activated non-catalytic subunit gamma 2 (*Prkag2*), phospholipase A2 (*Pla2*) and paraoxonase 2 (*Pon2*) in DSSR placentas was measured by quantitative PCR **(D)**. Human umbilical vein endothelial cell line EA.hy926 were incubated with the normal growth medium (basal), or additionally with serum (4 %) of control or citrulline-treated DSSR for 72 hours. Gene expression in the EA.hy926 incubated with the serum of either control or citrulline-treated rats was examined by qPCR **(E)**.*, p<0.05. Data are presented as mean ± SEM.

Placental gene expression of TNF-α and HIF-1α was downregulated by citrulline supplementation. Citrulline upregulated the gene expression of SIRT1, which is an important stress-response protein that modulates feto-placental vascular development ^44^ **(Figure 6C)**. The gene expression of enzymes related to lipid metabolism, including the paraoxonase 2 (PON2), protein kinase AMP-Activated non-catalytic subunit gamma 2 (Prkag2), phospholipase A2 (Pla2) was recently reported differentially expressed in placentas between Sprague Dawley (SD) rats and DSSR ^45^. Interestingly, the gene expression of PON2 and PLA2 was significantly upregulated in citrulline-treated DSSR **(Figure 6D)**.

In endothelial cell culture, incubation with the serum of control DSSR led to the upregulation of the TLR4 pathways (TLR4, MYD88 and p65) *in vitro*. These effects were largely normalized by the serum from citrulline-treated DSSR. Moreover, incubation with the serum of citrulline-treated DSSR significantly promoted the gene expression of PON2 **(Figure 6E)**. These results suggested that citrulline may regulate differential gene expression in DSSR placenta.

## Discussion

The present study demonstrates the protective effects of L-citrulline supplementation in an animal model of preeclampsia. Citrulline supplementation in DSSR leads to (i) reduction of maternal blood pressure and markers of preeclampsia; (ii) improvement of maternal endothelial function; (iii) amelioration of placental fibrosis and senescence; (iv) promotion of placenta angiogenesis; and (v) improvement of fetal growth.

### Significance of this study

Beneficial effects of citrulline supplementation on cardiometabolic health has been extensively studied ^46, 47^. However, the effect on preeclampsia and the long-term effects in gestation on fetal outcomes remain unclear. Clinical studies on citrulline levels in pregnancy have yielded conflicting results. One study have reported that citrulline can decrease blood pressure and improve vascular function in obese pregnant women ^48^. Another study found no correlation between citrulline level and blood pressure in severe preeclampsia ^49^. In a small-cohort of patients with severe preeclampsia (n=17), higher plasma citrulline levels were found comparing with controls or patients with mild or moderate preeclampsia ^50^. The authors interpreted the citrulline increase as a compensatory phenomenon. Maternal citrulline supplementation has been shown to enhance fetal growth and protein metabolism in a rat model of low protein diet-induced IUGR ^21, 51^. By far, there is no animal study investigating the effect of citrulline supplementation in preeclampsia. Citrulline supplementation is reported to be safe and well-tolerated as a single oral dose (2 - 15 g) in healthy adults ^52^. Indeed, citrulline has a limited degradation in the placenta, and can be efficiently transferred from the mother to the fetus to facilitate fetal growth and development ^53^. These make citrulline a good and safe candidate for targeting preeclampsia. The present study has presented the beneficial effects of citrulline supplementation in a rat model of superimposed preeclampsia. Multiple mechanisms could be involved in the beneficial effects of citrulline in preeclampsia, including (i) maternal vascular function, (ii) improved placental function, and (iii) a direct effect on the fetus.

### Maternal vascular function

In the endothelium, eNOS is important as it is responsible for the biosynthesis of NO from arginine, which confers vasorelaxing and antiplatelet properties ^54^. During pregnancy, the demand for arginine is increased due to the placental and embryonic growth. This demand can exceed the endogenous *de novo* synthesis of arginine from citrulline ^15^. NO production is increased during normal pregnancy and decreases in preeclampsia ^55, 56^. Arginine concentrations are significantly reduced in patients with preeclampsia when compared with healthy women ^57^. Reduction in eNOS/NO exacerbates sFlt-1-assoicated preeclampsia-like phenotype in mice ^58^. NOS inhibition by L-NAME perfusion has been used in rodent models to mimic preeclampsia-like phenotype ^59–61^. These suggest that NO plays a crucial role in maintaining a health pregnancy. Many studies have demonstrated vascular and endothelial dysfunction in preeclampsia ^62–65^. Vascular dysfunction in preeclampsia can be presented as augmented arterial stiffness and reduced vasodilatation ^65^.

Citrulline and watermelon extract supplementations have been demonstrated to reduce blood pressure in both pre-hypertensive and hypertensive patients ^66–69^. Citrulline can improve endothelial vasodilator function and reduce resting blood pressure by increasing the synthesis of arginine and NO levels ^70^. On the other hand, citrulline has been shown to promote the adaptations of the blood vessels to physiological and environmental stressors and improve blood flow responses by reducing wall stiffness ^46^. Our data suggests that citrulline supplementation can improve maternal vascular function in preeclampsia, possible *via* increasing eNOS expression and the production of NO from the endothelium in aorta. Blood pressure in the pregnant DSSR may be also reduced by the improved EDHF-mediated vasorelaxation in the resistance arteries. In addition, Maternal citrulline therapy has been shown to prevent L-NAME-induced programmed hypertension in rat ^71^, and the beneficial effect of citrulline to restore of the disturbed reactive oxygen species (ROS)/NO balance is proposed.

### Placental inflammation and fibrosis

Poor placentation leads to the early onset of preeclampsia. Preeclampsia can occur in patients with hydatidiform moles ^72^, which indicates that the placenta itself, but not the fetus, is required for the development of preeclampsia. Currently, the only effective treatment for preeclampsia is the delivery of the infant and placenta. NO participates in the process of placentation, placental angiogenesis and endothelial function in the placenta ^73^. Recent studies have suggested the importance of NO signaling in modulating the angiogenic factors including PlGF, VEGF, angiopoietin (Angpt), thrombospondin (Thbs) and anti-angiogenic factors like sFlt1 ^74^. The disturbance of the expression of these factors is associated with placental insufficiency and preeclampsia ^75^. Citrulline may improve placental function and fetal growth by promoting NO synthesis. Our data suggests a promotion of placental angiogenesis in the placentas of DSSR, which is associated with an upregulation of angiogenic factors and a downregulation of anti-angiogenic factor.

In preeclampsia, oxidative stress plays a pivotal role in the decreased NO bioavailability. Citrulline has antioxidant effects which can be NO-dependent or NO-independent. As mentioned, citrulline increases eNOS expression and activity, which in turn promotes NO production and reduces the formation of ROS. Decreased level of PON2 has been shown in DSSR and women with preeclampsia ^76^. Our data suggests that citrulline supplementation can upregulate the expression of PON2, which has protective role against oxidative stress and is associated with lipid metabolism.

Comparing to another study using a rat model of IUGR induced by maternal dietary protein restriction ^51^, we coherently observed no significant change in placental eNOS gene expression by citrulline supplementation. Opposite to their finding, we observed a downregulation of placental iNOS, which could be due to the inhibition of NF-κB-mediated transcription of iNOS in pregnant DSSR by citrulline. NF-κB-mediated inflammatory cytokines and oxidative stress may exert a pronounced effect in the fetus. Blunted upregulation of iNOS is suggested to attenuate preeclampsia ^77^. Citrulline can downregulate the expression of NF-κB and its downstream inflammatory markers in the placentas of DSSR. In addition, citrulline supplementation has been shown to selectively reduce the production of pro-inflammatory cytokines (such as IL-6, TNF-α, and C-reactive protein) ^78^, while preserving the production of anti-inflammatory cytokines (IL-10) ^79^.

TGF-β mediated fibrosis is one of the most prominent pathologic features of preeclamptic placentas ^39^. DSSR has increased placental hypoxia, which is a known factor that leads to fibrosis ^80^. Although the mechanism of fibrosis in the preeclamptic placentas is largely unknown, our results show that citrulline can reduce placental fibrosis in DSSR, which is associated with the downregulation of HIF-1α. A significant upregulation of TLR4 signaling pathway in the placentas of DSSR compared to SD rats has been reported ^81^. The activation of TLR4 signaling pathway in the placenta promotes pro-inflammatory cytokines production through the upregulation of several transcription factors, including NF-κB, which can induce inflammatory responses and placental cytotrophoblast apoptosis ^82^. Inhibition of HIF-1α-mediated TLR4 activation can attenuate apoptosis and promote placental angiogenesis during severe preeclampsia ^83^. Our results suggest that citrulline supplementation improves placental phenotype in DSSR, at least partly, by downregulating HIF-1α-mediated TLR4 activation in the placentas.

### Placental senescence

Premature ageing of the placenta has been recently associated with pregnancy complications including preeclampsia and IUGR ^40^. Preeclamptic women has increased placental senescence compared to normal pregnant women ^40^. Protein and gene expression of senescence markers (p16, p21, p53) are upregulated in the placenta from preeclamptic patients ^84, 85^. Chronic low-grade inflammation, increased placental oxidative stress and endoplasmic reticulum stress can facilitate placental senescence ^86^. DSSR has increased placental hypoxia.

NO bioavailability is reduced in senescent cells, while increasing NO bioavailability can activate telomerase and delay endothelial cell senescence ^87^. Citrulline may retard endothelial senescence by rescuing NO levels. Our results demonstrate that L-citrulline can also attenuate placental senescence in preeclampsia. LKB1 is highly expressed in senescent cells and can cause vascular senescence ^41^ and inhibit VEGF-induced angiogenesis ^42^. LKB1 is also an upstream inducer of AMP-activated protein kinase (AMPK) and amino acid-transporter required for mammalian target of rapamycin complex 1 (mTORC1) activity, which are upregulated in preeclampsia ^88^. However, the detailed involvement of LKB1 in preeclampsia remains unclear. The reduction of LKB1 expression in the placentas of citrulline-treat DSSR can be attributed to the increased expression of placental SIRT1. Indeed, SIRT1 is an important regulator of LKB1 level ^89^. Reduced expression and activity of SIRT1 in endothelial cells leads to the accumulation of acetylated LKB1, which cause senescence ^41, 89^. Therefore, citrulline may attenuate placental senescence in preeclampsia *via* activating SIRT1-mediated downregulation of LKB1. Further studies are warranted to explore the detailed mechanism of placental senescence in which may provide novel therapeutic strategies for preeclampsia.

### Lipid metabolism

Apart from the beneficial effects in improving vascular function and blood flow, citrulline supplementation may affect the efficiency of nutrient transfer in placentas. Consistent with another study ^90^, we observed an increased the fetal-to-placental weight ratio, an index of placental efficiency, by citrulline supplementation. This suggests that citrulline can improve the placental structural and functional adaptations to maintain fetal growth.

In a recent temporal transcriptomic analysis study, many differentially expressed genes between SD rats and DSSR are reported ^45^. Among these genes, we have noticed a cluster of genes which are genes involved in lipid and fatty acid metabolism, including the apolipoprotein family, *Pon2, Prkag2* and Pla2. PON2 is an important antioxidant enzyme that is associated with lipid metabolism and insulin sensitivity ^91, 92^. PLA2 is involved in the regulation of fatty acid and phospholipid metabolism while Prkag2 is involved in glucose and lipid metabolism ^93^. Lipids and fatty acids supplied through the placenta are important for fetal growth. However, excessive flux of lipids could promote oxidative stress and inflammatory cascades, contributing to preeclampsia ^94^. Increased level of circulating proinflammatory cytokines in preeclampsia are known to mediate lipid dysregulation. These result in augmented levels of circulating triglycerides (TG) and non-esterified free fatty acids (FFA) ^95^. However, the association between lipid metabolism and preeclampsia remains largely unclear. As forementioned, LKB1 is the upstream inducer of AMPK. Hyperactivation of AMPK is found in the placentas from preeclamptic patients, which is associated with disturbed fatty acid metabolome and impaired trophoblast invasion ^96^. In the present study, we report an upregulation of *Pon2* and *Pla2* in the placentas of citrulline-treated DSSR. As mentioned, our results show the upregulation of placental SIRT1 by citrulline. The expression of SIRT1, an nutrient sensor, is downregulated in preeclamptic women in comparison to healthy pregnant ^97^. The SIRT1-peroxisome proliferator-activated receptor gamma (PPARγ) axis is proposed to be a key player in trophoblast differentiation and placental development ^44^. The downregulation of SIRT1 expression in the placenta might be responsible for the increased NF-κB transcriptional activity and reduction of PPARγ ^98^. PPARγ is a transcription factor that is well-known for its key role in glucose and lipid metabolism ^44^. These results suggest an important link between placental lipid metabolism and preeclampsia, which warrants further studies.

### Limitation of the study

In the present study, we have shown that maternal citrulline treatment can improves preeclampsia pathologies and fetal growth. Nevertheless, we do not know the exact molecular mechanisms of citrulline yet. Our results have shown that the incubation of serum from citrulline-DSSR can ameliorate placental senescence and regulate the expression of different genes in endothelial cells *in vitro*. This suggests the possible contribute of citrulline-related metabolites in the beneficial effects of citrulline supplementation. The possible mechanism may involve NO-dependent and/or NO-independent pathways. It is well established that citrulline can be converted into arginine thereby improving NO production. NO may mediate transcriptional regulation of histone-modifying enzymes through the formation of S-nitrosothiols or iron nitrosyl complexes ^99^. Metabolites of citrulline and arginine, such as polyamines, are required in multiple stages of pregnancy, including implantation, early embryogenesis, fetal growth, and placental development ^100^. Polyamine metabolism plays an important role in placental function. Reductions in polyamine bioavailability in pregnant rodents have been associated with abnormal placentation and fetal growth restriction ^101, 102^. Recent study also reveals the epigenetic effect of placental polyamines by regulating acetyl-CoA level and histone acetylation ^100^. Thus, citrulline metabolites may contributes to the effects observed in the present study, although we did not examine the metabolite changes in the placentas of DSSR with citrulline supplementation. Another possible mechanism is a direct stimulation of protein synthesis to improve fetal growth ^103^, probably by activating the phosphorylation of proteins in the mTOR signaling pathway ^19^.

A second limitation is that we did not examine the reprogramming effect of maternal citrulline supplementation (i.e., the disease risk of the offspring at adult age). Such a reprogramming effect has been reported for citrulline in the spontaneously hypertensive rats. Maternal treatment with citrulline of the spontaneously hypertensive rats ameliorated the development of hypertension in the offspring ^104^. We have observed an improvement of fetal growth in the DSSR. Thus, a reprogramming effect of maternal citrulline treatment is conceivable in the DSSR. Nevertheless, this was not investigated, because it is out of scope of this study. This cardiovascular and metabolic disease risk of the offspring from citrulline-treated mothers should be further addressed in future studies.

In conclusion, this study shows that L-citrulline supplementation in a rat model of superimposed preeclampsia can reduce gestational hypertension, improve placentation and fetal growth. L-citrulline could be a potent and safe therapeutic strategy for preeclampsia that benefit both the mother and the fetus.

### Perspective

There is currently no effective pharmacotherapy to treat preeclampsia. The effect of maternal citrulline supplementation in preeclampsia and the long-term effects in gestation on fetal outcomes remain unclear. Results from our study indicate that L-citrulline supplementation could represent a new therapeutic strategy. Further research studies will be focused on the effect of L-citrulline in nutrient metabolism of the placenta and fetus, as well as the reprogramming effect of L-citrulline in the growth and development of the offspring. Promising results have been achieved, although only limited relevant clinical studies are currently available. We expect the results presented in this study to provide a strong scientific basis for citrulline supplementation to improve maternal health and placental development in preeclampsia. L-citrulline could be a potent and safe therapeutic strategy for preeclampsia that benefit both the mother and the fetus. Clinical trials would be warranted to explore the beneficial effects of L-citrulline in human preeclampsia.

## Author contributions

NX and HL designed the study. AWCM, YZ, UDPL, GR, AW and AH performed the experiments and analyzed data. AWCM wrote the manuscript. AD, TM, EC, NX and HL critically reviewed and edited the manuscript. All authors agreed to its publication.

