## Supplementary file for "L-citrulline ameliorates pathophysiology in a rat model of superimposed preeclampsia"

**Supplementary table 1.** Primer list for gene expression studies using qPCR

| | |
|---|---|
| Rat-TGFb-F | CTGCTGACCCCCACTGATAC |
| Rat-TGFb-R | AGCCCTGTATTCCGTCTCCT |
| Rat-mmp2-F | CAAGCCCAAGTGGGACAAGA |
| Rat-mmp2-R | CCATGCTCCCATCGACCAAA |
| Rat-mmp9-F | AGCCGACGTCACTGTAACTG |
| Rat-mmp9-R | AACAGGCTGTACCCTTGGTC |
| Rat-PlGF-F | CAGCCAACATCACTATGCAG |
| Rat-PlGF-R | TCCTCTGAGTGGCTGGTTA |
| Rat-VEGF-F | GCAATGATGAAGCCCTGGAG |
| Rat-VEGF-R | GGTGAGGTTTGATCCGCATG |
| Rat-FGF2-F | TCCATCAAGGGAGTGTGTGC |
| Rat-FGF2-R | TCCGTGACCGGTAAGTGTTG |
| Rat-flt-1-F | GTACCTCACCGTGCAAGGAA |
| Rat-flt-1-R | ACTTCGGAAGAAGACCGCTT |
| Rat-thbs-1-F | AAGACGTCGACGAGTGCAAA |
| Rat-thbs-1-R | CAGGCAGTTGTAGCCAGGAT |
| Rat-thbs-2-F | CGTGAGCGATGAGAAGGACA |
| Rat-thbs-2-R | CGATCTGTGCTTGGTTGTGC |
| Rat-angpt2-F | CATGATGTCATCGCCCGACT |
| Rat-angpt2-R | TCCATGTCACAGTAGGCCTTG |
| Rat-LKB1-F | GGCCAACGTCAAGAAGGAGA |
| Rat-LKB1-R | AGTACCCATGAGCTTGGCAC |
| Rat-p16-F | TCGTACCCCGATACAGGTGAT |
| Rat-p16-R | TGTCTAGGAAGCCCTCCCG |
| Rat-p21-F | GACAGGGTGTTGACGATGCC |
| Rat-p21-R | TGTGCCTTAAGAAAGAGTACAAACT |
| Rat-p53-F | CCCCTGAAGACTGGATAACTGT |
| Rat-p53-R | AACTCTGCAACATCCTGGGG |
| Rat-TLR4-F | ATGCCTCTCTTGCATCTGGC |
| Rat-TLR4-R | ATTGTCTCAATTTCACACCTGGA |

| | |
|---|---|
| Rat-MYD88-F | GCAGTTTGTTGGATGCCTGG |
| Rat-MYD88-R | TTCTGGCAGTCCTCCTCGAT |
| Rat-p65-F | AGCCATAGCCATAGTTGCGG |
| Rat-p65-R | CGTCATCCTTAAGGGCCTGAT |
| Rat-p50-F | GCTTACGGTGGGATTGCATT |
| Rat-p50-R | TTATGGTGCCATGGGTGATG |
| Rat-iNOS-F | TCCTCAGGCTTGGGTCTTGT |
| Rat-iNOS-R | AGAAACTTCCAGGGGCAAGC |
| Rat-VCAM-1-F | GAAGCCGGTCATGGTCAAGT |
| Rat-VCAM-1-R | GGTCACCCTTGAACAGTTCTATCTC |
| Rat-ICAM-1-F | CGGGAGATGAATGGTACCTACAA |
| Rat-ICAM-1-R | TGCACGTCCCTGGTGATACTC |
| Rat-HIFa-F | CAACTGCCACCACTGATGAA |
| Rat-HIFa-R | TGGGTAGAAGGTGGAGATGC |
| Rat-TNFa-F | GGGGCCACCACGCTCTTCTGTC |
| Rat-TNFa-R | TGGGCTACGGGCTTGTCACTCG |
| Rat-SIRT1-F | TTCCTGTGGGATACCTGACTTCA |
| Rat-SIRT1-R | TGCCTTGAGGATCTGGGAGAT |
| Rat-pla2-F | GATCACCTGCAGCGACAAAA |
| Rat-pla2-R | GGGGTGACAGCCTAACAGTG |
| Rat-prkag2-F | CCATGCTGATCCGTGTCCT |
| Rat-prkag2-R | CACTCTCTGAGTCTTCTTCCTCC |
| Rat-Pon2-F | TTCCAAACTGCCGCCTCATT |
| Rat-Pon2-R | AAACTTGAGGCCCACGCTAA |
| Human-TLR4-F | AAAATCCCCGACAACCTCCC |
| Human-TLR4-R | TGTCTGGATTTCACACCTGGA |
| Human-MYD88-F | GACCCAGCATTGAGGAGGAT |
| Human-MYD88-R | CTGCACAAACTGGATGTCGC |
| Human-p65-F | GCTGCATCCACAGTTTCCAG |
| Human-p65-R | TCCCCACGCTGCTCTTCTAT |
| Human-ICAM-1-F | TCCCCCCGGTATGAGATTG |

| | |
|---|---|
| Human-ICAM-1-R | GCCTGCAGTGCCCATTATG |
| Human-PON2-F | GGGTAGGCTGTCATCCTAATGG |
| Human- PON2-R | TGTAGGCTTCTCAGATAGAATGTTCTG |
| Human-p53-F | GCGAGCACTGCCCAACA |
| Human-p53-R | TGAAATATTCTCCATCCAGTGGTTT |
| Human-eNOS-F | CACCAGGAAGAAGACCTTTAAAGAA |
| Human-eNOS-R | TCACTCGCTTCGCCATCAC |
| Human-SIRT1-F | TCAGTGGCTGGAACAGTGAG |
| Human-SIRT1-R | AGCGCCATGGAAAATGTAAC |
| Human-IL6-F | CCGGGAACGAAAGAGAAGCT |
| Human-IL6-R | GCGCTTGTGGAGAAGGAGTT |
| Human-NF-kB-F | CCCTGAGACAAATGGGCTACAC |
| Human-NF-kB-R | CAGCGAGTGGGCCTGAGA |
| Human-TNFa-F | GGTTTGCTACAACATGGGCTACA |
| Human-TNFa-R | TGGCCCAGGCAGTCAGA |
| Human-HIFa-F | TGCTCATCAGTTGCCACTTC |
| Human-HIFa-R | TCCTCACACGCAAATAGCTG |
| Human-thbs-1-F | AAGACGCCTGCCCCATCAAT |
| Human-thbs-1-R | CTGTACCCCTCCTCCACAGG |
| Human-bactin-F | CCTGGCACCCAGCACAAT |
| Human-bactin-R | GCCGATCCACACGGAGTACT |
| Rat-GAPDH-F | TTCTTGTGCAGTGCCAGCC |
| Rat-GAPDH-R | CGTCCGATACGGCCAAATC |
